# Formation of periodic pigment spots by the reaction-diffusion mechanism

**DOI:** 10.1101/403600

**Authors:** Baoqing Ding, Erin L. Patterson, Srinidhi V. Holalu, Jingjian Li, Grace A. Johnson, Lauren E. Stanley, Anna B. Greenlee, Foen Peng, H. D. Bradshaw, Benjamin K. Blackman, Yao-Wu Yuan

## Abstract

Many organisms exhibit visually striking spotted or striped pigmentation patterns. Turing’s reaction-diffusion model postulates that such periodic pigmentation patterns form when a local autocatalytic feedback loop and a long-range inhibitory feedback loop interact. At its simplest, this network only requires one self-activating activator that also activates a repressor, which inhibits the activator and diffuses to neighboring cells. However, the molecular activators and repressors fully fitting this versatile model remain elusive. Here, we characterize an R2R3-MYB activator and an R3-MYB repressor in monkeyflowers that correspond to Turing’s model and explain how periodic anthocyanin spots form. Notably, disrupting this pattern impacts pollinator visitation. Thus, subtle changes in simple reaction-diffusion networks are likely essential contributors to the evolution of the remarkable diversity of periodic pigmentation patterns in flowers.

## Maintext

Periodic pigmentation patterns like the stripes of zebras, the spiral pigmentation of seashells, and the petal spots of many flowers have fascinated biologists and mathematicians for centuries. One proposed developmental explanation for how such periodic patterns form is Turing’s reaction-diffusion model (*1*), in which dynamic and autonomous patterns are generated simply owing to the interaction of an activator and a repressor. The activator self-activates and activates the repressor, which then diffuses and inhibits the activator along the diffusion path. This mechanism amplifies initial cellular fluctuations into tissue-level spatial patterns (*2-4*). Computer simulations suggest that by tinkering with the diffusion constants and the kinetics of the activator-repressor interaction, this simple circuit can recapitulate the immense diversity of pigmentation patterns observed in nature (*3*). However, the molecular identities and dynamics of actual activator-repressor pairs that fulfill the classic Turing model for pigment patterning have remained elusive. Anthocyanin spots in flower petals provide an excellent empirical system to reveal the molecular basis for the formation and evolution of periodic pigmentation patterns. These patterns, which are highly diverse in the angiosperms even among different varieties of the same species (*5, 6*), are known to serve as critical cues in plant-pollinator interactions (*7-9*); and the genetic network controlling anthocyanin pigment production is otherwise well described (*10, 11*).

Previously, we identified a self-activating R2R3-MYB transcription factor, NECTAR GUIDE ANTHOCYANIN (NEGAN), that positively regulates anthocyanin spot formation in the monkeyflower species *Mimulus lewisii* (Fig. 1A) and hypothesized that a mechanism akin to the Turing model could generate this periodic pattern (*12*). If so, a corresponding repressor of NEGAN function must exist. While down-regulation of *NEGAN* expression abolishes anthocyanin spot formation in the nectar guides (i.e., the two yellow ridges of the ventral petal; Fig. 1B), loss-of-function mutation in the presumptive repressor gene should cause expansion of the spots. To search for such a repressor, we carried out an ethyl methanesulfonate mutagenesis screen using the *M. lewisii* inbred line LF10 and isolated one candidate mutant, named *red tongue* (*rto*), that fits the expectation. Homozygous *rto* mutants develop a near uniform patch of anthocyanin color across the entire nectar guide region (Fig. 1C). Coincidentally, we discovered naturally occurring mutants with *rto*-like phenotypes segregating in several geographically distinct populations of the congener *M. guttatus* (fig. S1), and we genetically examined the two variants found at Sweetwater Creek (SWC) in Oregon (Fig. 1F) and Littlerock Dam (LRD) in Southern California (Fig. 1G). A complementation cross indicated that these natural *M. guttatus* variants represent different alleles of the same locus (fig. S1B). The *rto* mutant and *rto-*like natural alleles behave semi-dominantly; F_1_s derived by crossing these alleles to corresponding wild-type individuals exhibit an intermediate phenotype (Fig. 1D,F,G).

**Fig. 1.**
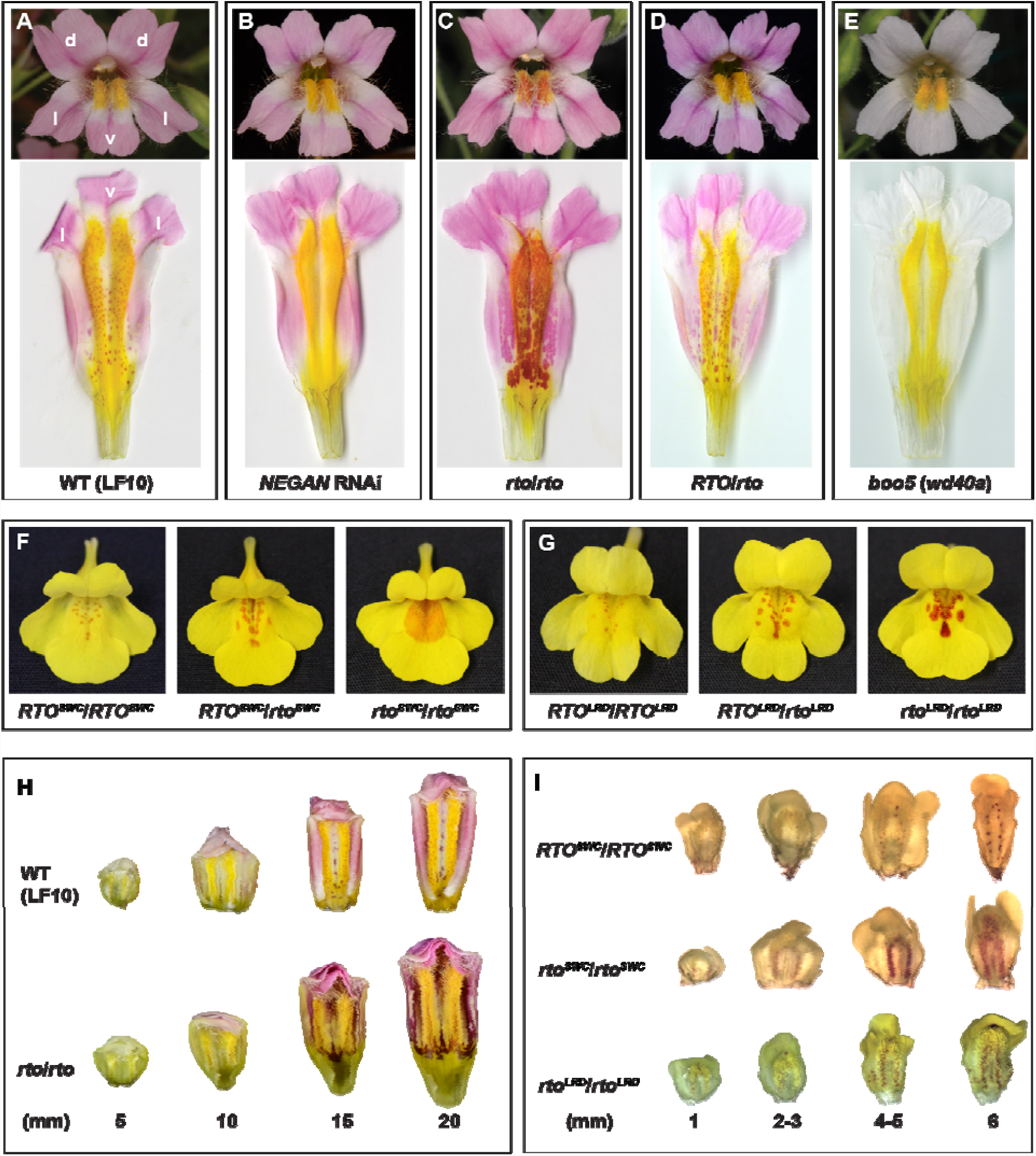
Periodic anthocyanin spots in *Mimulus lewisii* and *M. guttatus*. (**A-E**) Anthocyanin spots on the yellow background of the wild-type (LF10) and various mutants and transgenic lines. d: dorsal; l: lateral; v: ventral. (**F-G**) Anthocyanin spot patterns of natural variants of *M. guttatus* segregating in the SWC and LRD populations. (**H-I**) Corresponding developmental series of anthocyanin spot formation in *M. lewisii* and *M. guttatus*.

Although *M. lewisii* and *M. guttatus* diverged about 20 million years ago (*13*), the similarity and specificity of the altered pigmentation phenotypes led us to suspect that all these mutant phenotypes are caused by lesions in the same gene. To identify the gene responsible for the *rto* phenotype in each species, we performed bulked segregant analyses on the *M. lewisii* mutant and the two *rto-*like *M. guttatus* alleles (i.e., *rto*^*SWC*^ and *rto*^*LRD*^). As expected, the independent mapping experiments all pinpointed the same causal genomic interval (Fig. 2A-C). Inspection of the bulked segregant reads aligning to this ~60 kb region in the wild-type *M. lewisii* (LF10) genome revealed only one mutation, which causes an aspartic acid-to-glycine replacement at a highly conserved site of a small R3-MYB protein (the ortholog of Migut.B01849) that is 91 amino acids long (Fig. 2D and fig. S2A). Furthermore, fine mapping resolved the *M. guttatus rto*^*SWC*^ allele to an interval spanning four genes (fig. S2B), including the same *R3-MYB* gene *Migut.B01849*. Sanger sequencing of this gene revealed a 7-bp insertion in the second exon, which alters the reading frame, and a premature stop mutation in the third exon of *M. guttatus* SWC and LRD *rto*-like plants, respectively (Fig. 2D and fig. S2A). Notably, this R3-MYB is closely related to a group of R3-MYBs that are known to repress anthocyanin biosynthesis, including ROSE INTENSITY1 (ROI1) in *M. lewisii* (*14*), MYBx in *Petunia* (*15*), and CAPRICE in *Arabidopsis* (*16*) (fig. S2A). Taken together, these results strongly suggest this *R3-MYB* as the causal gene underlying the *RTO* locus in both *M. lewisii* and *M. guttatus*.

**Fig. 2.**
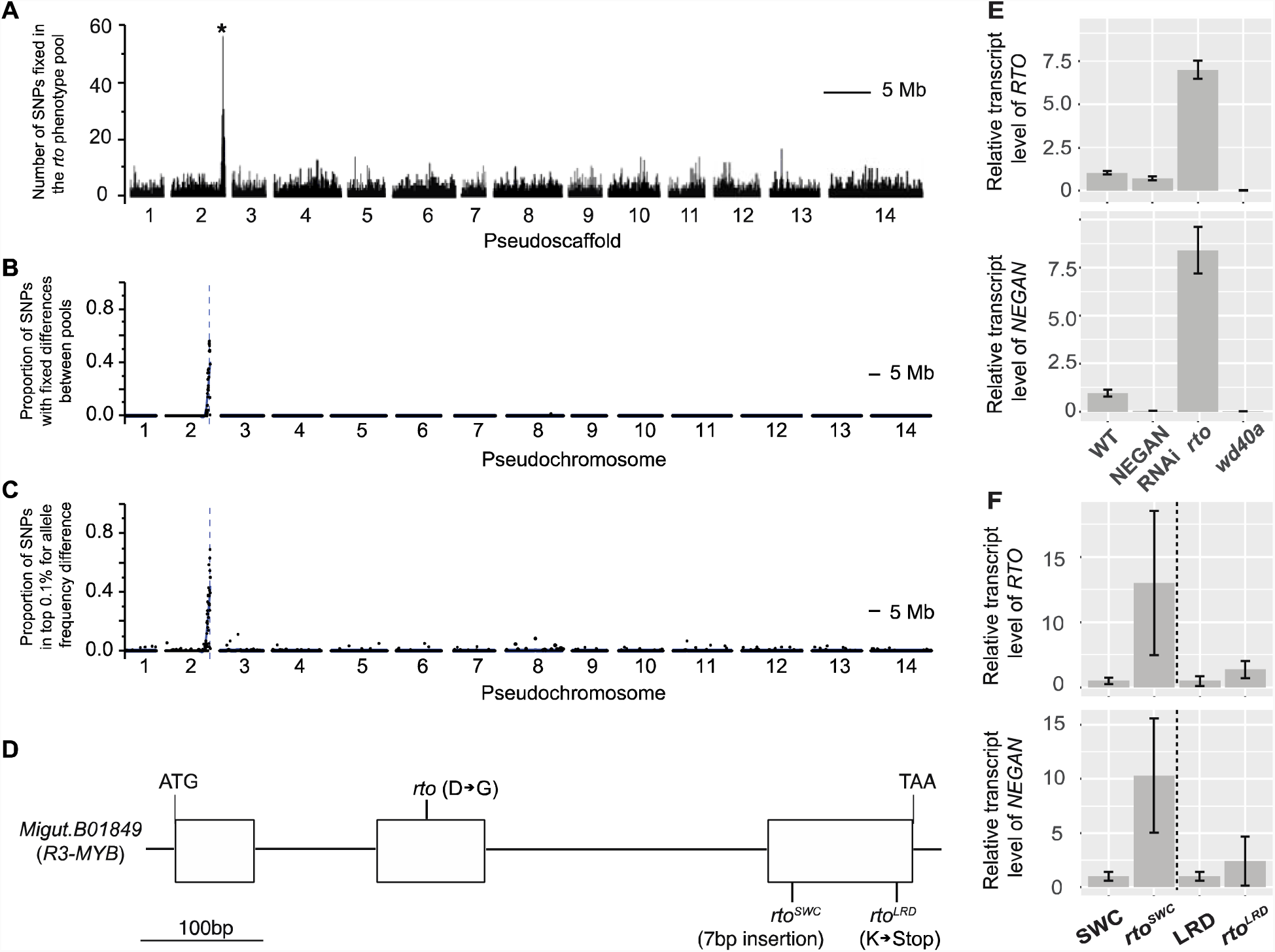
Identification of the *RTO* gene in *Mimulus lewisii* and *M. guttatus* and its relative expression in various mutant and transgenic lines. (**A-C**) Bulked segregant analyses of *rto* (A), *rto*^*SWC*^ (B), and *rto*^*LRD*^ (C) narrowed *RTO* down to the same genomic interval. (**D**) Schematic of the *RTO* (*R3-MYB*) gene, showing the molecular lesions of the three mutant alleles. (**E-F**) Relative transcript level of *RTO* (upper) and *NEGAN* (lower) in *M. lewisii* (E) and *M. guttatus* (F) as measured by qRT-PCR, standardized to the corresponding wild-type (LF10 for *M. lewisii*, SWC or LRD for *M. guttatus*). Dashed lines separate two different experiments. Error bars represent 1 SD from three biological replicates.

To verify the function of this *R3-MYB* gene, we knocked down its expression level in *M. lewisii* and *M. guttatus* by RNA interference (RNAi). We obtained multiple independent transgenic lines from both species with severe phenotypes that completely recapitulate the mutant phenotypes (Fig. 3A and B). Some lines in the *M. lewisii* background have an even stronger phenotype than *rto* (Fig. 3A), suggesting that *rto* is a hypomorphic instead of null allele. Notably, CRISPR/Cas9-mediated frameshift mutations in *MgRTO* also recapitulate the *rto* phenotype (Fig. 3B and fig. S3). Together, the association of expanded pigmentation with three independent lesions in *RTO* and with RNAi knockdown and/or targeted disruptions of *RTO* in two congeneric species strongly indicates that *RTO* functions as a repressor of anthocyanin spot patterning. As such, we expected that over-expression of *RTO* from the CaMV *35S* promoter would attenuate or completely abolish the anthocyanin spots. As expected, the corollas of strong *35S:RTO* lines in *M. lewisii* are completely white and lack anthocyanin spots in the nectar guide region (Fig. 3C), confirming that *RTO* acts as a potent repressor of anthocyanin pigmentation.

**Fig. 3.**
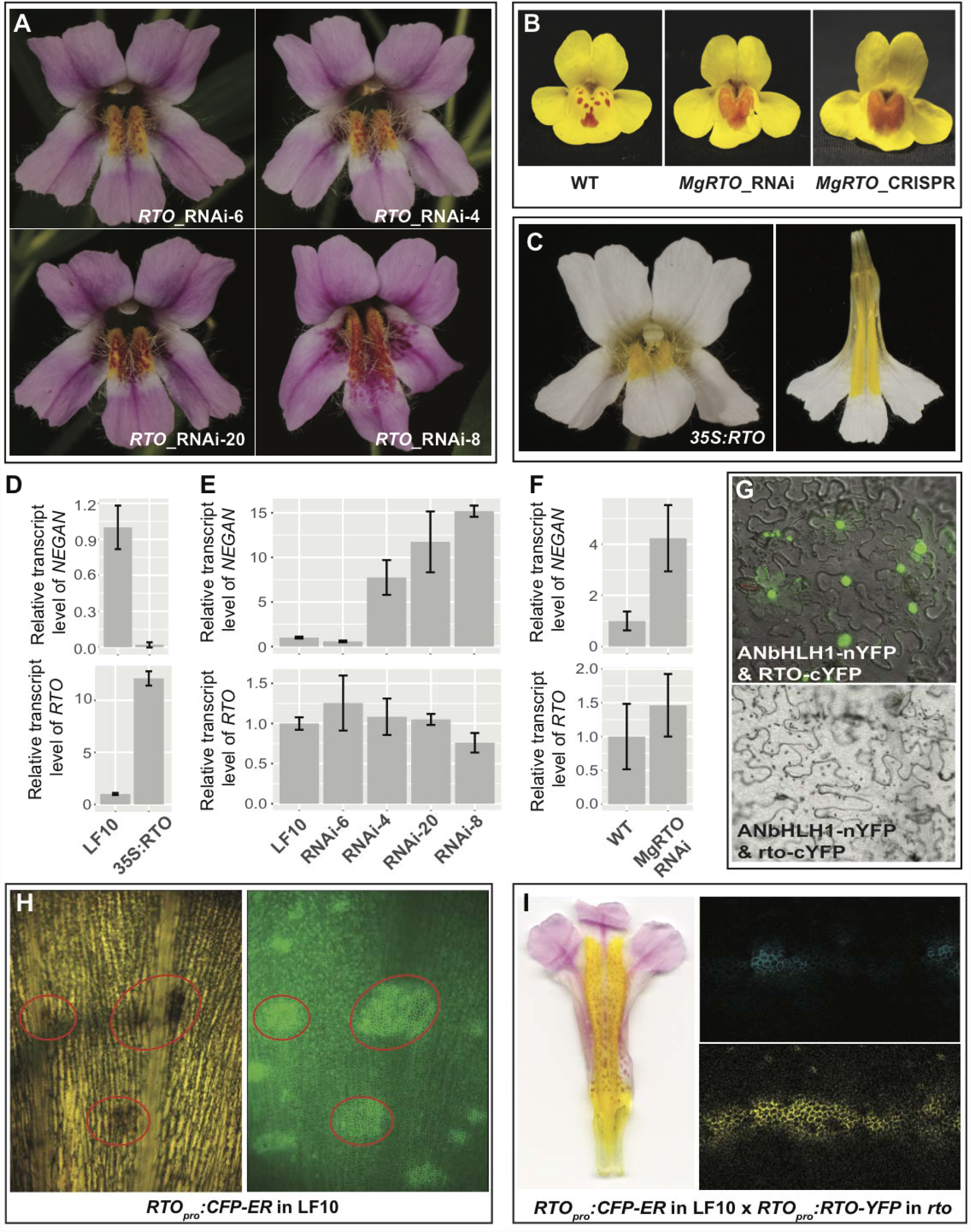
Molecular characterization of the NEGAN-RTO interaction. (**A**) RNAi of *RTO* in *Mimulus lewisii* generates a range of anthocyanin spot patterns. (**B**) RNAi and CRISPR-mediated genome editing of *RTO* in *M. guttatus* recapitulates the *rto*-like phenotype. (**C-D**) Over-expression of *RTO* in *M. lewisii* abolishes anthocyanin production throughout the corolla (C) and causes dramatic down-regulation of *NEGAN* (D). (**E-F**) Relative expression of *NEGAN* across *RTO* RNAi lines is positively correlated with anthocyanin production in the nectar guides, but *RTO* transcript level shows no correlation to the strength of the phenotype in *M. lewisii* (E) and *M. guttatus* (F). All relative transcript levels are measured by qRT-PCR, standardized to the corresponding wild-type. Error bars represent 1 SD from three biological replicates. (**G**) BiFC assay shows that the wild-type RTO protein interacts with ANbHLH1, whereas the mutant rto protein does not. (**H**) Spatial pattern of *RTO* promoter activity revealed by the *RTO*_*pro*_:*CFP-ER* construct. CFP fluorescence signal foreshadows and co-localizes with anthocyanin production (red ovals). (**I**) F_1_ hybrids between a *RTO*_*pro*_:*CFP-ER* line (in the wild-type background) and a *RTO*_*pro*_:*YFP-RTO* line (in the *rto* background) bear flowers similar in anthocyanin phenotype to the non-transgenic *RTO/rto* heterozygote and reveal a broader spatial distribution of RTO protein (yellow) than *RTO* promoter activity (blue).

Having identified the putative repressor, we next tested the other predictions of the classic Turing model: (i) the self-activating activator (i.e., NEGAN) also activates the repressor (i.e., RTO); (ii) the repressor inhibits the activity of the activator; and (iii) the repressor “diffuses” from cell to cell.

To test the first prediction that the activator also activates the repressor, we compared *RTO* gene expression levels between wild type and *NEGAN* RNAi plants as well as the *M. lewisii wd40a* mutant. The *wd40a* mutant is relevant because in all angiosperms characterized to date, the anthocyanin-activating R2R3-MYB transcription factor (e.g., NEGAN) functions in an obligatory protein complex that also contains a bHLH protein and a WD40 protein (*10, 11*). In *M. lewisii* flowers, the bHLH and WD40 components of the complex (MlANbHLH1 and MlWD40a, respectively) are expressed in the entire corolla, while the R2R3-MYB component is supplied by the complementary tissue-specific expression of two paralogs, *NEGAN* and *PETAL LOBE ANTHOCYANIN* (*PELAN*), that are expressed in nectar guides and petal lobes, respectively (*12*). The *wd40a* mutant completely lack anthocyanins in both their petal lobes and the nectar guides (ref. *12*; Fig. 1E). Down-regulation of *NEGAN* by RNAi abolishes the anthocyanin spots in the nectar guides but does not affect the petal lobe color (Fig. 1B), whereas the loss-of-function *pelan* mutant and *PELAN* RNAi lines lose all the anthocyanin pigments in the petal lobe but with unaffected nectar guide spots (ref. *12*; fig. S4A). If nectar guide expression of *RTO* is activated by NEGAN acting in a complex with MlANbHLH1 and MlWD40a, we would expect reduced *RTO* expression in *NEGAN* RNAi lines and the *wd40a* mutant compared to wild-type plants.

Indeed, for whole corollas sampled from 10 mm stage buds, the developmental stage when both *NEGAN* and *RTO* show peak expression level and the anthocyanin spots just start appearing (Fig. 1H and fig. S4B), the *RTO* transcript level decreased to an undetectable level in the *wd40a* mutant compared to the wild-type (Fig. 2E). *NEGAN* expression is also abolished in the *wd40a* mutant (Fig. 2E), as previously shown (*12*), because the ability of *NEGAN* to self-activate also requires the complete NEGAN-MlANbHLH1-MlWD40a protein complex. In the 10-mm stage whole corollas of *M. lewisii NEGAN* RNAi lines, however, the *RTO* transcript level only slightly decreased compared to the wild-type (Fig. 2E). We predicted this finding resulted from activation of *RTO* expression in the petal lobe by *PELAN*. Indeed, when examined in dissected petal lobes and nectar guides at this stage, petal lobe *RTO* expression is unaffected in the *NEGAN* RNAi lines, but is reduced to undetectable level in the *pelan* mutant; whereas nectar guide *RTO* expression is unaffected in the *pelan* mutant, but clearly reduced in the *NEGAN* RNAi lines (fig. S4C). These results suggest that *RTO* expression is activated by PELAN and NEGAN in the petal lobe and nectar guides, respectively, and requires the same bHLH and WD40 components in both tissues. That *PELAN* cannot self-activate (*12*) explains the generally uniform anthocyanin pigmentation in the petal lobes, compared to the periodic spots in the nectar guides. Similarly, when *NEGAN* expression was knocked down in *M. guttatus*, the anthocyanin spots completely disappeared from nectar guides of strong RNAi lines and *RTO* expression levels decreased accordingly (fig. S5). Taken together, our results confirmed the first prediction; the activator, *NEGAN*, activates the expression of the repressor, *RTO*, in the nectar guides of both *Mimulus* species.

To test the second prediction that the repressor inhibits the activator function, we assayed relative *NEGAN* expression levels in the *rto* mutant and *rto*-like natural variants as well as various *RTO* transgenic lines. Since *NEGAN* expression is self-activated (*12*), if RTO inhibits the activity of NEGAN, loss of *RTO* function should result in higher *NEGAN* expression level and over-expression of *RTO* should have the opposite effect. These are precisely the patterns we observed: *NEGAN* transcript level increased 6-8-fold in *rto* and *rto*^*SWC*^ (Fig. 2E,F) and decreased >10-fold in the strong 35S:*RTO* lines (Fig. 3C,D). The increase of *NEGAN* transcript level in *rto*^*LRD*^ is less dramatic (Fig. 2F), indicating that *rto*^*LRD*^ is a weak allele, consistent with the phenotype (Fig. 1G). In the *M. lewisii RTO* RNAi lines, *NEGAN* expression is up-regulated up to 15-fold and there is a positive correlation between *NEGAN* expression level and the severity of the anthocyanin phenotype (Fig. 3A and D). Likewise, *MgNEGAN* expression is also up-regulated in the *MgRTO* RNAi lines (Fig. 3F). These results strongly indicate that the repressor (RTO) does inhibit the activity of the activator (NEGAN).

That the transcript level of *RTO* itself in the *RTO* RNAi lines is not lower than in the wild-type (Fig. 3E, F) seems surprising at first glance. However, this finding is consistent with the anticipated dynamic interactions between *NEGAN* and *RTO*: as shown above, knock-down of *RTO* transcript levels by RNAi resulted in up-regulation of *NEGAN*, which in turn would up-regulate *RTO*, but the increased *RTO* transcript level would then be counterbalanced by RNAi. Consequently, unlike *NEGAN, RTO* transcript level showed no clear correlation to the severity of the anthocyanin phenotype in the *RTO* RNAi lines.

The mechanism by which R3-MYB repressors inhibit R2R3-MYB activator function is well understood for their homologous proteins in *Arabidopsis* and *Petunia* (*15, 16*). These R3-MYB proteins have neither DNA-binding nor activation domains, but they compete with the R2R3-MYB for the limited supply of the bHLH proteins, sequestering the bHLH proteins into inactive complexes. Like NEGAN, RTO also contains the bHLH-interacting motif in its R3 domain (*17*). The aspartic acid-to-glycine replacement in the mutant rto protein disrupts the bHLH-interacting motif (fig. S2A), and therefore, is expected to attenuate or abolish its interaction with MlANbHLH1. Bimolecular fluorescence complementation assays confirmed this hypothesis. The wild-type RTO protein interacts with MlANbHLH1, whereas the mutant rto protein does not (Fig. 3G).

To test the third prediction that the repressor “diffuses” to neighboring cells (i.e., intercellular movement) –– we focused on RTO because homologous R3-MYBs have been shown to move between cells (*15, 18*) whereas the corresponding R2R3-MYB activators function cell autonomously (*15, 19*). One way to test the mobility of RTO is to compare the spatial pattern of *RTO* transcription with the RTO protein distribution (*18*). If RTO moves between cells, RTO proteins should be observable over a broader domain of cells than where *RTO* is transcribed.

To make this comparison, we first examined the spatial pattern of *RTO* transcription by transforming the wild-type *M. lewisii* (LF10) with a construct that expresses cyan fluorescent protein (CFP) with an endoplasmid reticulum (ER) retention signal, driven by the *RTO* promoter (~1.7kb DNA fragment upstream of the translation initiation site). The CFP signal marks the cells with *RTO* promoter activity, as the CFP-ER product cannot move from the source cell to neighboring cells. Multiple independent *RTO*_*pro*_:*CFP-ER* lines showed the same pattern; CFP fluorescence signal co-localizes with the anthocyanin spots (Fig. 3H), consistent with the expectation that both *RTO* and the anthocyanin biosynthetic genes are regulated by the same transcriptional activator (i.e., NEGAN).

Next, to reveal the spatial distribution of RTO proteins, we expressed a YFP-RTO fusion protein driven by the same *RTO* promoter in the *rto* background. In the majority of the *RTO*_*pro*_:*YFP-RTO* transformants, the nectar guide anthocyanin pigmentation was reduced (fig. S6A), and the extent of this “rescue” of the *rto* phenotype was positively correlated with the transgene expression level (fig. S6B). Thus, the YFP-RTO fusion protein retains the function of RTO as an anthocyanin repressor *in vivo*. The complete elimination of nectar guide anthocyanin rather than restoration to the wild-type pattern for a substantial number of *RTO*_*pro*_:*YFP-RTO* lines in the *rto* background was unexpected (fig. S6A). One possible explanation is that a recessive epistatic interaction between the hypomorphic *rto* allele and the transgene shifts the dynamics of the NEGAN-RTO interaction. However, when we crossed representative *RTOpro:CFP-ER* and *RTOpro:YFP-RTO* lines to bring the two transgenes together in a *RTO/rto* heterozygous background, we observe typical *RTO/rto* anthocyanin patterning rather than the “over-rescue” phenotype (Figs. 1D and 3I), indicating wild-type NEGAN-RTO dynamics are re-established. In this background, YFP-RTO signal was detected not only in the source cells of the *RTO* promoter activity, as reflected by the CFP-ER signal, but also in neighboring cells (Fig. 3I and fig. S7). Taking these transcript and protein localization results together, we conclude that the *RTO* gene is transcribed in the anthocyanin spot cells but the RTO protein moves to adjacent cells.

To gain further insight into the mechanism of RTO movement, we generated *35S:YFP-RTO* transformants in *M. lewisii*. Like the strong *35S:RTO* lines, strong *35S:YFP-RTO* transgenic lines produce completely white flowers, confirming again that the YFP-tag does not interfere with RTO function. We found YFP-RTO localized to both nucleus and cytoplasm. We also found many punctate YFP-RTO signals, mainly in the cytoplasm. These puncta travel from the nucleus to membrane (fig. S8; Movie S1), reminiscent of CAPRICE, which is known to interact with protein transport partners and travel through the endosome system in *Arabidopsis* roots (*20*). To determine if the structures surrounding YFP-RTO are endocytic membrane components, we tested whether YFP-RTO co-localizes with the fluorescence signal of FM4-64, which is known to stain cellular membranes and endocytic structures (*21*). Indeed, YFP-RTO and FM4-64 showed perfect co-localization (fig. S9). These results suggest that inter-cellular RTO movement is likely to be mediated by the endosome system.

Having demonstrated that NEGAN and RTO function as a activator-repressor pair that fits the Turing model in generating periodic anthocyanin spots, we next explored how variants that alter the NEGAN-RTO interaction dynamics are distributed in natural populations of *M. guttatus* (fig. S1) and their potential impacts on plant performance. In two of three population surveys for SWC and LRD, we found that *M. guttatus rto-*like alleles segregate at low but non-trivial frequencies in the wild (2011 survey: *rto*^*SWC*^ frequency = 0.1, n=10 maternal individuals; 2017 surveys: *rto*^*LRD*^ frequency = 0.02, n=51 and *rto*^*SWC*^ frequency = 0, n=49). Moreover, *rto-*like plants have been seen in at least three additional populations, each >200km from either SWC or LRD (fig. S1). In one of these populations, *rto*-like plants carry an 8-bp deletion that removes a splice acceptor site and causes a frameshift at the start of the second exon of *RTO* (fig. S1), and we estimate this allele is present at a frequency of ~0.11 in the population based on phenotype (2018 survey: n=200 individuals).

The moderately high frequency of *rto-*like alleles in some *M. guttatus* populations and the fact that these anthocyanin spots serve as important nectar guides in the closely related *M. luteus* (*7*) motivated us to test whether natural variation in the anthocyanin spot patterns could impact pollinator visitation. We carried out a series of pollinator preference experiments in a laboratory setting using the *rto*^*SWC*^ allele since the heterozygous and homozygous genotypes form a continuum of spot expansion (Fig. 1F). In a direct comparison between genotypes, naïve pollinators preferentially visited *rto*^*SWC*^ homozygote flowers over wild-type flowers, and pollinators also preferred heterozygote to wild-type flowers (Tables 1, S1-S3). Pollinators did not discriminate between heterozygote flowers and *rto*^*SWC*^ homozygote flowers, however. Given that corolla UV pigmentation and cell shape do not differ between full-sib *RTO*^*SWC*^ homozygous vs. *rto*^*SWC*^ carrier flowers (Fig. S10), we conclude that altered anthocyanin pigmentation pattern is responsible for preferred visitation to *rto*^*SWC*^ homozygotes and *RTO*^*SWC*^*/rto*^*SWC*^ heterozygotes. Determining whether the *rto*-like alleles represent nascent selective sweeps or are polymorphisms maintained by frequency-dependent selection or at mutation-selection balance due to costs (e.g., florivore attraction) will require further study in field conditions. Nonetheless, these findings demonstrate the principle that changes in nectar guide patterning that arise by modulating the underlying reaction-diffusion system can influence fundamental plant-pollinator interactions.

**Table 1.** Anthocyanin spot patterning affects pollinator visitation. Pooled goodness-of-fit test over ten trials per pairwise *RTO* genotype combination. Significant heterogeneity G-values indicate variation in bee preference among individual trials; nonetheless, in all cases, the effects of genotype in significant individual trials were in the same direction as the pooled test (Tables S1-S3).

| Genotype | Observed Visits | Pooled G-value | d.f. | *P* | Heterogeneity G-value | d.f. | *P* |
| --- | --- | --- | --- | --- | --- | --- | --- |
| *RTO/RTO* | 246 | 30.99 | 1 | <0.001 | 83.34 | 9 | <0.001 |
| *rto/rto* | 393 |  |  |  |  |  |  |
| *RTO/RTO* | 450 | 17.92 | 1 | <0.001 | 37.01 | 9 | <0.001 |
| *RTO/rto* | 566 |  |  |  |  |  |  |
| *RTO/rto* | 404 | 1.09 | 1 | 0.3 | 14.18 | 9 | 0.12 |
| *rto/rto* | 420 |  |  |  |  |  |  |

In summary, we have described a simple reaction-diffusion network responsible for the formation of periodic anthocyanin spots in *Mimulus* flowers. This activator-repressor pair appears to fit the classical Turing model precisely. The activator, NEGAN, is self-activating and also activates the expression of the repressor, RTO; RTO inhibits the activity of NEGAN; and RTO shows intercellular movement. An array of anthocyanin spot patterns is achievable by modulating the dynamics of the activator-repressor interaction through experimental manipulations or natural mutations, and altering the spot patterns can impact pollinator visitation. Thus, subtle changes in this simple interacting network are likely essential contributors to the remarkable diversity of flower pigmentation patterns, and we expect that molecular circuits with similar properties do explain periodic pigmentation patterns in other systems as first envisioned by Turing more than six decades ago.

## Acknowledgements

We thank J. Manson for guidance with the pollinator cage experiments; S. Criswell and B. Glover for assistance and advice with SEM; C. O’Connell for assistance with confocal imaging; E. LoPresti and K. Toll for alerting us to natural *rto*-like variants; A. Nguyen, J. Weger, and S. McDevitt for assistance with library construction and sequencing; S. Lewis for assistance in EMS mutant screen, and T. Rushton, I. Tucker, I. Tan, W. Crannage, and the University of Connecticut, UC Berkeley Oxford Tract Facility, Duke University, and University of Washington research greenhouse staff for plant husbandry and phenotyping assistance; the Blackman Lab for feedback on the manuscript; and J. Willis for his generous support.

## Author contributions

Y.W.Y, B.K.B., and H.D.B. conceptualized and guided the project. B.D., J.L., L.E.S., and F.P. conducted the experiments in *M. lewisii*. E.L.P., S.H., G.J., A.B.G., and B.K.B. conducted the experiments in *M. guttatus*. B.D., E.L.P., S.H., B.K.B, and Y.W.Y. analyzed the data and wrote the manuscript.

## Data and materials availability

Genome sequence data have been deposited in the NCBI Sequence Read Archive (BioProject PRJNA481753). Requests for *M. lewisii* and *M. guttatus* materials should be made to Y.W.Y. and B.K.B., respectively.

## List of Supplementary Materials

Materials and Methods

Fig S1 - S10

Table S1 – S6

References (22 - 32)

Movie S1

## References and Notes

1. A. M. Turing. Philos. Trans. Royal Soc. B. 237, 37–72 (1952).

2. H. Meinhardt, A. Gierer. Bioessays 22, 753–760 (2000).

3. S. Kondo, T. Miura. Science 329, 1616–1620 (2010).

4. J. B. Green, J. Sharpe. Development 142, 1203–1211 (2015).

5. C.-C. Hsu, Y.-Y. Chen, W.-C. Tsai, W.-H. Chen, H.-H. Chen. Plant Physiol. 168, 175– 191(2015).

6. M. Yamagishi. Sci. Hort. 163, 27–36 (2013).

7. R. Medel, C. Botto-Mahan, M. Kalin-Arroyo. Ecology 84, 1721–1732 (2003).

8. Z.-X. Ren, D.-Z. Li, P. Bernhardt, H. Wang. Proc. Natl. Acad. Sci. U.S.A. 108, 7478– 7480 (2011).

9. M. L. de Jager, E. Willis-Jones, S. Critchley, B. J. Glover. Evol. Ecol. 31, 193–204 (2017).

10. K. M. Davies, N. W. Albert, K. E. Schwinn. Funct. Plant Biol. 39, 619–638 (2012).

11. B. J. Glover. Understanding Flowers and Flowering: An Integrated Approach. (Oxford University Press, Oxford, ed. 2nd, 2014).

12. Y.-W. Yuan, J. M. Sagawa, L. Frost, J. P. Vela, H. D. Bradshaw Jr. New Phytol. 204, 1013–1027 (2014).

13. Z. L. Nie, H. Sun, P. M. Beardsley, R. G. Olmstead, J. Wen. Am. J. Bot. 93, 1343–1356 (2006).

14. Y.-W. Yuan, J. M. Sagawa, R. C. Young, B. J. Christensen, H. D. Bradshaw. Genetics 194, 255–263 (2013).

15. N. W. Albert, K. M. Davies, D. H. Lewis, H. B. Zhang, M. Montefiori et al. Plant Cell 26, 962–980 (2014).

16. H. F. Zhu, K. Fitzsimmons, A. Khandelwal, R. G. Kranz. Mol. Plant. 2, 790–802 (2009).

17. I. M. Zimmermann, M. A. Heim, B. Weisshaar, J. F. Uhrig. Plant J. 40, 22–34 (2004).

18. T. Kurata, T. Ishida, C. Kawabata-Awai, M. Noguchi, S. Hattori et al. Development 132, 5387–5398 (2005).

19. J.-Y. Kim, Y. Rim, J. Wang, D. Jackson. Genes Dev. 19, 788–793 (2005).

20. K. Koizumi, S. Wu, A. MacRae-Crerar, K. L. Gallagher. Curr. Biol. 21, 1559–1564 (2011).

21. S. Bolte, C. Talbot, Y. Boutte, O. Catrice, N. Read, B. Satiat-Jeunemaitre. J. Microsc. 214, 159–173 (2004).

22. C. R. Owen, H. D. Bradshaw. Arthropod-Plant Interact. 5, 235–244 (2011).

23. Y.-W. Yuan, J. M. Sagawa, V. S. Di Stilio, H. D. Bradshaw. Genetics 194, 523–528 (2013).

24. A. J. Kelly, J. H. Willis. Mol. Ecol. 7, 769–774 (1998).

25. H. Li, R. Durbin. Bioinformatics 25, 1754–1760 (2009).

26. H. Li, B. Handsaker, A. Wysoker, T. Fennell, J. Ruan, N. Homer, G. Marth, G. Abecasis, R. Durbin. Bioinformatics 25, 2078–2079 (2009).

27. A. Kerschen, C. A. Napoli, R. A. Jorgensen, A. E. Muller. FEBS Lett. 566, 223–228 (2004).

28. M. Michniewicz, E. M. Frick, L. C. Strader. BMC research notes 8, 63 (2015).

29. K. W. Earley, J. R. Haag, O. Pontes, K. Opper, T. Juehne, K. M. Song, C. S. Pikaard. Plant J. 45, 616–629 (2006).

30. C. Grefen, N. Donald, K. Hashimoto, J. Kudla, K. Schumacher, M. R. Blatt. Plant J. 64, 355–365 (2010).

31. B. Ding, Y.-W. Yuan. Plant Cell Rep. 35, 771–777 (2016).

32. J. H. McDonald. Handbook of biological statistics. (Sparky House Publishing, Baltimore, 2014). pp. 53-58.

